# Reactivation-coupled brain stimulation enables complete learning generalization

**DOI:** 10.1101/2025.09.27.678935

**Authors:** Yibo Xie, Minmin Wang, Yuan Gao, Baoyu Wu, Shaomin Zhang, Mengyuan Gong, Zoe Kourtzi, Ke Jia

**Author notes:** Corresponding author: Dr. Ke Jia, MOE Frontier Science Center for Brain Science and Brain-machine Integration, Zhejiang University.

## Abstract

Generalization of learned knowledge to new contexts is essential for adaptive behavior. Despite extensive research on the brain plasticity mechanisms underlying learning specificity, the mechanisms that facilitate generalization remain poorly understood. Here, we investigate whether using brain stimulation to disrupt offline consolidation in visual cortex promotes learning generalization. Separate groups of participants (N = 144) were trained on visual detection tasks using either a reactivation-based protocol or conventional full-practice, combined with anodal or sham transcranial direct current stimulation (tDCS) over the visual cortex. Strikingly, only combination of reactivation-based learning with anodal tDCS produced complete generalization from trained to untrained stimuli, an effect consistently replicated across features (orientation, motion direction). In contrast, reactivation-based learning alone and conventional full-practice – whether with or without brain stimulation – yielded stimulus-specific learning. Importantly, reactivation-coupled brain stimulation achieved generalization with an 80% reduction in training trials while maintaining learning gains comparable to full-practice. These findings demonstrate that reactivation and neuromodulation interact to unlock learning generalization, revealing a key brain plasticity mechanism and offering a rapid, translatable strategy for sensory rehabilitation.

## Introduction

Extensive practice over days to months can yield highly specialized skills, but the hallmark of learning is generalization – an ability to transfer acquired skills flexibly to new contexts (Shepard, 1987). Previous research has manipulated behavioral protocols to identify factors that shape the degree of specificity and transfer, including task difficulty (Ahissar and Hochstein, 1997; Jeter et al., 2009), training duration (Jeter et al., 2010), stimulus (Yashar and Denison, 2017) and task variability (Manenti et al., 2023; Xiao et al., 2008). These strategies can promote generalization, but often entail trade-offs: protocols that are simpler or shorter tend to yield smaller learning gains, whereas those incorporating stimulus or task variability often require training durations comparable to or exceeding that of conventional full-practice regimens. Moreover, although such behavioral manipulations reveal empirical benefits, they provide limited insight into the underlying neural mechanisms, leaving a critical gap in our understanding of how brain plasticity supports the generalization of learning.

To address this gap, we focus on visual perceptual learning (VPL), a well-established model of experience-dependent improvements in perceptual decisions (Watanabe and Sasaki, 2015). A hallmark of VPL is its high degree of stimulus specificity, a phenomenon thought to reflect over-specialized neural representations in the visual cortex (i.e., perceptual overfitting) (Sagi, 2011). This overfitting may arise due to learning either modifying feature representation in early visual cortex (Jia et al., 2020, Jia et al., 2024; Yan et al., 2014) or enhancing read-out of sensory neurons from early visual areas to optimize perceptual decisions (Dosher and Lu, 2017; Law and Gold, 2008), with greater specificity emerging as training progresses. This creates a central paradox in which extensive training is required to achieve substantial learning gains, yet such training simultaneously drives overfitting that limits the generalizability of learning.

While improvements in conventional VPL primarily depend on prolonged online practice, recent studies propose an alternative mechanism based on offline memory consolidation, which may help resolve this paradox. Specifically, reactivation-based protocol uses brief reminder trials to retrieve existing perceptual memories and enables learning via offline consolidation processes. Although behavioral improvements – both in overall learning gains and in specificity – are comparable between full-practice and reactivation-based VPL, evidence indicates that the latter engages distinct brain plasticity processes. Specifically, these learning gains are thought to arise from offline stabilization mediated by γ-aminobutyric acid (GABA) (Eisenstein et al., 2023), the brain’s primary inhibitory neurotransmitter. Anodal transcranial direct current stimulation (tDCS), a non-invasive brain stimulation technique, has been shown to reduce GABA concentrations in the visual cortex (Barron et al., 2016). Therefore, we hypothesize that applying anodal tDCS to the visual cortex may disrupt perceptual overfitting in reactivation-based VPL, thereby enhancing the generalization of learned perceptual skills.

To test this hypothesis, we trained separate groups of participants on visual detection tasks using either reactivation-based or full-practice protocols, combined with anodal or sham tDCS over the visual cortex. Only combination of reactivation-based learning and anodal tDCS produced complete transfer of learning from trained to untrained stimuli, an effect consistently replicated across stimulus features (orientation, motion direction). In contrast, reactivation alone or full-practice protocols resulted in stimulus-specific learning. These findings reveal a key brain plasticity mechanism enabling generalization and suggesting a rapid, transferable training strategy with direct relevance for clinical rehabilitation (e.g. sensory deficits).

## Results

We trained forty-eight adults with a reactivation-based learning protocol (Bang et al., 2018), using an orientation detection task (Figure 1A-B). On each trial, participants viewed two sequentially presented stimuli and reported which interval contained the target (a Gabor patch embedded in noise). Participants were randomly assigned to either the Anodal or Sham group (N = 24 per group). The anodal electrode was placed over O1 (contralateral to the trained visual field) and the cathodal electrode over Cz (vertex), following the international 10-20 EEG system.

**Figure 1.**
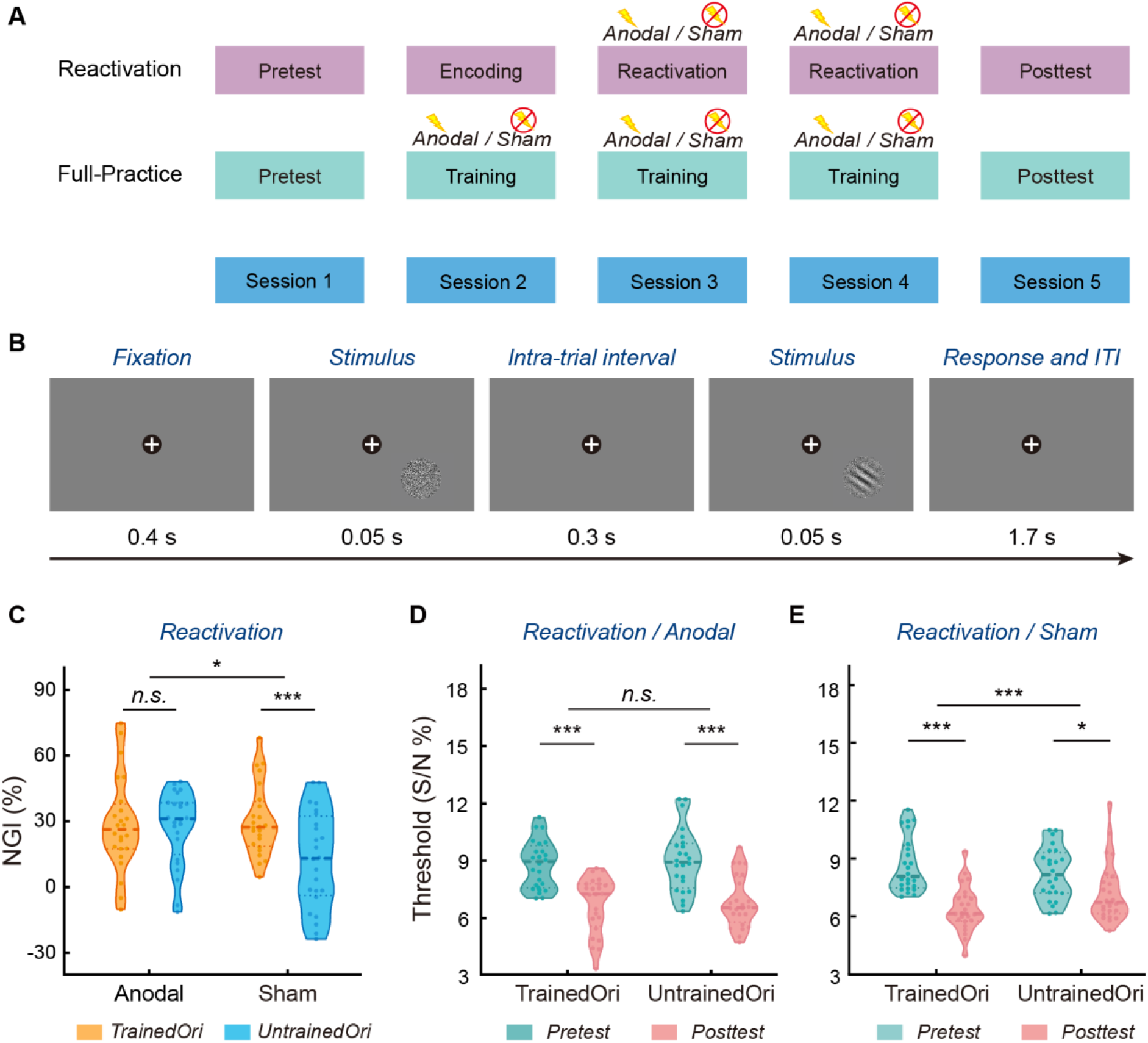
Procedures and transfer effect using orientation task. (A) Experimental design. Participants completed five sessions, including a pretest, training or reactivation, and a posttest. (B) Orientation detection task. Participants reported which of the two intervals contained the Gabor orientation. (C) Normalized learning gain index (NGI) for trained versus untrained orientations in the Reactivation (Anodal vs. Sham) groups. (D) Thresholds (S/N ratio) in *Reactivation/Anodal* group. A two-way repeated measures ANOVA (session: pretest vs. posttest × orientation: trained vs. untrained) revealed a significant main effect of session (F(1,23) = 124.406, *p* < 0.001, η^2^_p_ = 0.844), but no interaction effect (F(1,23) = 0.010, *p* = 0.923, BF_01_ = 3.633), demonstrating comparable learning for the trained (paired t-test: t(23) = 7.338, *p* < 0.001, Cohen’s d = 1.498) and untrained (paired t-test: t(23) = 7.765, *p* < 0.001, Cohen’s d = 1.585) orientations. (E) Thresholds (S/N ratio) in *Reactivation/Sham* group. A two-way repeated measures ANOVA (session: pretest vs. posttest × orientation: trained vs. untrained) revealed a significant interaction (F(1,23) = 16.477, *p* < 0.001, η^2^_p_ = 0.417), demonstrating stronger learning for the trained (paired t-test: t(23) = 8.386, *p* < 0.001, Cohen’s d = 1.712) compared to the untrained (paired t-test: t(23) = 2.599, *p* = 0.016, Cohen’s d = 0.531) orientation. The central lines in the box plot indicate the median values. The upper and lower lines represent the interquartile range (25^th^ – 75^th^ percentiles). Each dot represents data from one participant. ^*^*p* < 0.05, ^***^*p* < 0.001, *n*.*s*. = not significant.

Our results showed complete transfer of learning in the Reactivation/Anodal group. In contrast, we observed orientation-specific learning in the Reactivation/Sham group (Figure 1C), consistent with previous studies (Amar-Halpert et al., 2017). To quantify the learning effects, we calculated a normalized learning gain index (NGI = [(Pre-test threshold – Post-test threshold) / ((Pre-test threshold + Post-test threshold) / 2)] × 100 %). A two-way mixed ANOVA (stimulation condition: anodal vs. sham × orientation: trained vs. untrained) on NGI revealed a significant interaction (F(1,46) = 5.551, *p* = 0.023, *η*^*2*^_*p*_ = 0.108). Post-hoc comparisons showed significantly greater improvements for the trained over untrained orientation in the Reactivation/Sham group (paired t-test: t(23) = 4.484, *p* < 0.001, Cohen’s d = 0.915), but comparable improvements across orientations in the Reactivation/Anodal group (paired t-test: t(23) = 0.213, *p* = 0.833, BF_01_ = 4.563). Similar results were obtained when analyzing thresholds (i.e., S/N ratio, Figure 1D-E). These findings indicate that reactivation-coupled occipital stimulation promotes full transfer of learning.

To validate that the observed generalization was not specific to the orientation detection task, we conducted a follow-up experiment using motion detection (N = 48, Figure 2A), while keeping the task structure and stimulation protocols similar. A two-way mixed ANOVA (stimulation condition: anodal vs. sham × direction: trained vs. untrained) on NGI revealed a significant interaction (F(1,46) = 13.107, p < 0.001, *η*^*2*^_*p*_ = 0.222, Figure 2B). Post-hoc comparisons indicated that the Reactivation/Sham group exhibited significantly greater learning for the trained than untrained direction (paired t-test: t(23) = 4.285, p < 0.001, Cohen’s d = 0.875), whereas the Reactivation/Anodal group showed equivalent improvements across directions (paired t-test: t(23) = 0.117, p = 0.908, BF_01_ = 4.630). Similar results were obtained when analyzing motion coherence (Figure 2C-D). This replication with a different stimulus feature corrraborates our results showing that reactivation-coupled anodal stimulation enables complete transfer of perceptual learning.

**Figure 2.**
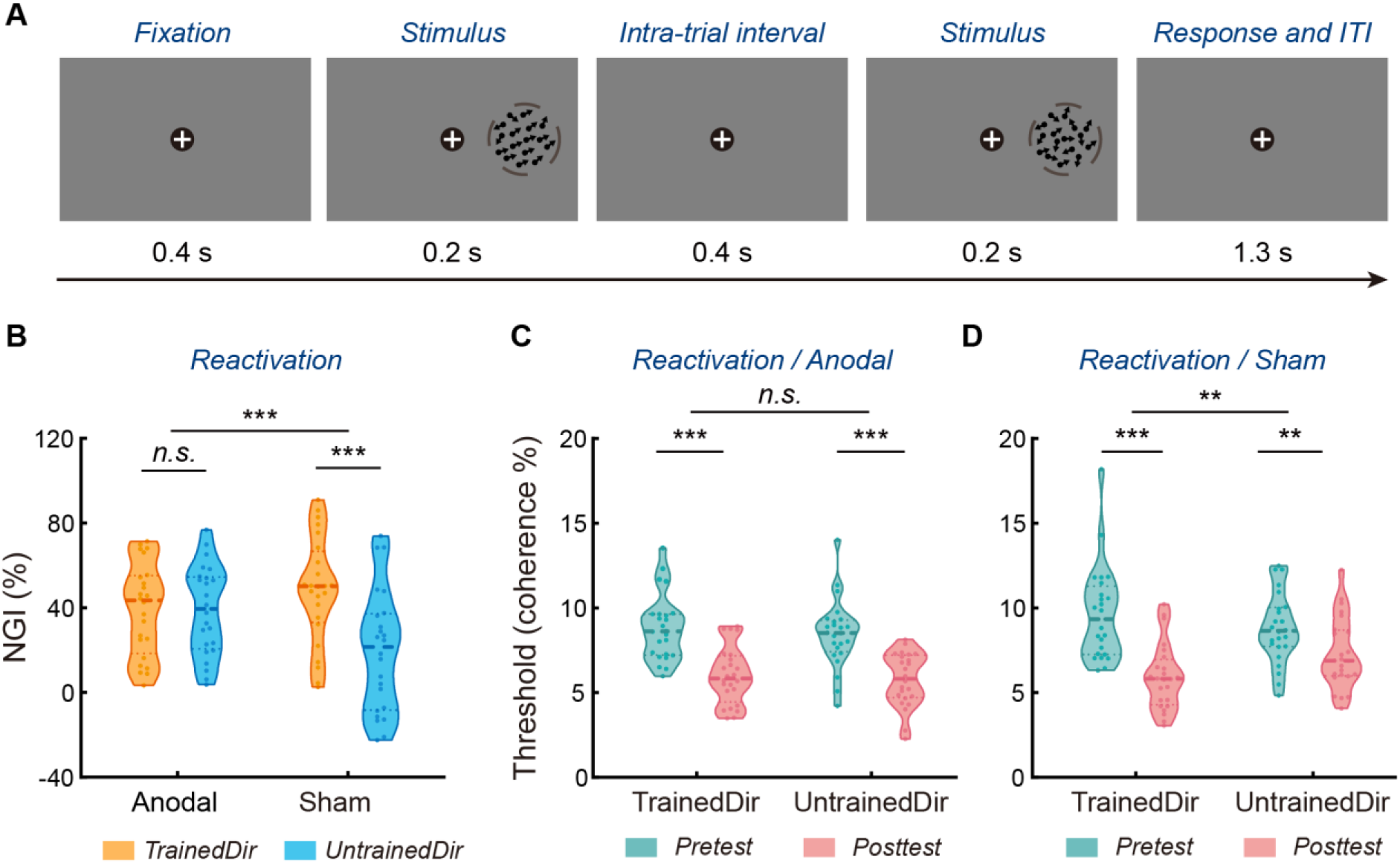
Transfer effect using motion detection task. (A) Motion detection task. Participants reported which of the two intervals contained the coherent motion dot field. (B) Normalized learning gain index (NGI) for the trained versus untrained direction in the Reactivation (Anodal vs. Sham) groups. (C) Thresholds (motion coherence) in *Reactivation/Anodal* group. A two-way repeated measures ANOVA (session: pretest vs. posttest × direction: trained vs. untrained) revealed a significant main effect of session (F(1,23) = 102.652, *p* < 0.001, η^2^_p_ = 0.817), but no interaction effect (F(1,23) = 0.252, *p* = 0.621, BF_01_ = 3.239), demonstrating comparable learning for the trained (paired t-test: t(23) = 9.134, *p* < 0.001, Cohen’s d = 1.864) and untrained (paired t-test: t(23) = 8.973, *p* < 0.001, Cohen’s d = 1.832) directions. (D) Thresholds (motion coherence) in *Reactivation/Sham* group. A two-way repeated measures ANOVA (session: pretest vs. posttest × orientation: trained vs. untrained) revealed a significant interaction (F(1,23) = 9.864, *p* = 0.005, η^2^_p_ = 0.300), demonstrating larger learning effect for the trained (paired t-test: t(23) = 7.764, *p* < 0.001, Cohen’s d = 1.585) compared to the untrained (paired t-test: t(23) = 3.244, *p* = 0.004, Cohen’s d = 0.662) direction. The central lines in the box plot indicate the median values. The upper and lower lines represent the interquartile range (25^th^ – 75^th^ percentiles). Each dot represents data from one participant. ^**^*p* < 0.01, ^***^*p* < 0.001, *n*.*s*. = not significant.

We next asked whether the observed generalization in Reactivation/Anodal group might reflect a reduced overall amount of learning. To test this possibility, we recruited an additional group of 24 adults who completed standard full-practice with sham stimulation on the orientation detection task (Figure 1A). A one-way ANOVA on NGI for the trained orientation (group: Reactivation/Anodal vs. Reactivation/Sham vs. Full-Practice/Sham) revealed no significant main effect of group (F(2,69) = 0.085, *p* = 0.919), which was further supported by a Bayesian analysis (BF_01_ = 7.952; see Materials and Methods). Direct comparison confirmed that the Reactivation/Anodal group attained learning gains comparable to those of the Full-Practice/Sham group (independent t-test: t(46) = 0.095, *p* = 0.925, BF_01_ = 3.467). These results suggest that the reactivation-coupled brain stimulation enhanced generalization without compromising overall learning gains.

Next, we examined whether the observed generalization in Reactivation/Anodal group could be attributed to reduced learning specificity inherent to the reactivation-based training protocol itself. To address this, we conducted a two-way mixed ANOVA (group: Reactivation/Sham vs. Full-practice/Sham × orientation: trained vs. untrained) on NGI. The analysis revealed a robust main effect of orientation (F(1,46) = 21.870, *p* < 0.001, *η*^*2*^_*p*_ = 0.322), but no significant effects of training protocol (F(1,46) = 0.029, *p* = 0.866, BF_01_ = 3.486) or their interaction (F(1,46) = 0.882, *p* = 0.352, BF_01_ = 2.302). These results indicate that both the Reactivation/Sham and Full-Practice/Sham groups showed a comparable degree of learning specificity. Taken together with the performance in Reactivation/Anodal group, these findings demonstrate that anodal tDCS is necessary for achieving enhanced generalization in reactivation-based VPL.

Further, anodal tDCS combined with full-practice failed to enhance generalization; instead, it increased learning specificity. We recruited another 24 adults and applied anodal tDCS in combination with full-practice. A two-way mixed ANOVA (group: Full-Practice/Anodal vs. Full-Practice/Sham × orientation: trained vs. untrained) on NGI revealed a significant interaction (F(1,46) = 4.237, *p* = 0.045, *η*^*2*^_*p*_ = 0.084, Figure 3), indicating enhanced specificity in the Full-Practice/Anodal group (i.e., larger improvements for the trained than untrained orientation). Post-hoc comparisons showed that this increased specificity was driven by greater gains for the trained orientation with anodal than sham tDCS (independent t-test: t(46) = 2.489, *p* = 0.017, Cohen’s d = 0.718), but no difference for the untrained orientation (independent t-test: t(46) = -0.246, *p* = 0.807, BF_01_ = 3.395). These results indicate that anodal tDCS combined with full-practice increased learning specificity, rather than enhancing generalization.

**Figure 3.**
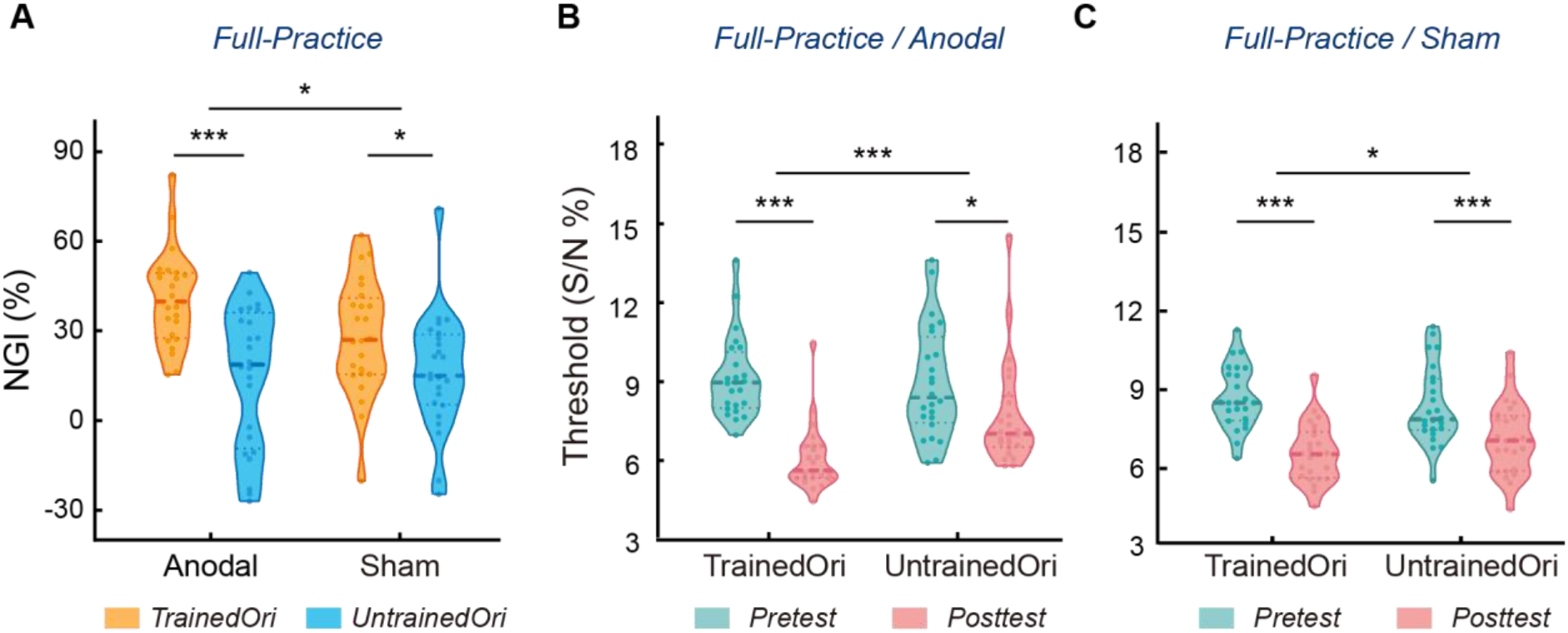
Full-Practice leads to learning specificity. (A) Normalized learning gain index (NGI) for the trained versus untrained orientation in the Full-Practice (Anodal vs. Sham) groups. (B) Thresholds (S/N ratio) in Full-Practice/Anodal group. A two-way repeated measures ANOVA (session: pretest vs. posttest × orientation: trained vs. untrained) revealed a significant interaction (F(1,23) = 17.961, p < 0.001, η^2^_p_ = 0.438), demonstrating larger learning effect for the trained (paired t-test: t(23) = 10.689, p < 0.001, Cohen’s d = 2.182) compared to the untrained (paired t-test: t(23) = 2.782, p = 0.011, Cohen’s d = 0.568) orientation. (C) Thresholds (S/N ratio) in Full-Practice/Sham group. A two-way repeated measures ANOVA (session: pretest vs. posttest × orientation: trained vs. untrained) revealed a significant interaction (F(1,23) = 5.380, p = 0.030, η^2^_p_ = 0.190), demonstrating larger learning effect for the trained (paired t-test: t(23) = 6.932, p < 0.001, Cohen’s d = 1.415) compared to the untrained (paired t-test: t(23) = 4.101, p < 0.001, Cohen’s d = 0.837) orientation. The central lines in the box plot indicate the median values. The upper and lower lines represent the interquartile range (25th – 75th percentiles). Each dot represents data from one participant. ^*^p < 0.05, ^***^p < 0.001, n.s. = not significant.

Lastly, to statistically validate the enhanced generalization in the Reactivation/Anodal group compared with other groups, we calculated the NGI difference between the trained and untrained orientation (NGI difference = NGI_trained_ – NGI_untrained_), where lower values indicate greater transfer and higher values indicate greater specificity. A two-way independent-measures ANOVA (training protocol: reactivation vs. full-practice × stimulation condition: anodal vs. sham) on NGI difference showed a significant interaction (F(1,92) = 9.755, *p* = 0.002, *η*^*2*^_*p*_ = 0.096). This pattern showed that Reactivation/Anodal enhanced generalization, whereas Full-Practice/Anodal increased specificity. The divergence across protocols suggests distinct neural mechanisms for reactivation-based versus repetition-based learning, consistent with previous studies (Eisenstein et al., 2023; Kondat et al., 2024).

## Discussion

Our study provides the first evidence that reactivation and neuromodulation interact to unlock complete learning generalization. Reactivation-based protocols have been shown to result mainly in learning specificity (Amar-Halpert et al., 2017), with a recent report suggesting partial transfer (Kondat et al., 2025), consistent with the effects observed for the sham condition in our study. In contrast, we demonstrate that reactivation-coupled anodal tDCS enables complete learning transfer. We have replicated this result across two perceptual detection tasks with different stimulus features (orientation, motion direction), providing evidence for a perceptual plasticity mechanism that boosts generalization. In particular, our protocol may disrupt the offline consolidation of learning in visual cortex – likely due to tDCS-mediated GABA reduction – thereby reducing perceptual overfitting and promoting generalization. This reduction in overfitting does not compromise learning gains, in line with previous work suggesting that reactivation-based plasticity involves higher-order areas (Kondat et al., 2024).

Previous studies on repetition-based learning have shown that different forms of electrical stimulation can modulate learning outcomes in distinct ways. For instance, anodal or cathodal stimulation alters VPL in a task-specific manner (Frangou et al., 2018), while stimulation at alpha – but not theta or gamma – frequencies can enhance VPL improvements (He et al., 2022). Consistent with these findings, the present study further demonstrated that anodal tDCS selectively enhanced stimulus-specific learning. This pattern of results, that is distinct from that observed in the reactivation groups, may be explained by two factors. First, prior research suggests that excessive training in full-practice group can induce hyper-stabilization of memory traces in the visual cortex (Shibata et al., 2017), a process that may rely less on offline consolidation. As a result, the GABA reduction induced by anodal tDCS exterted a diminished impact on consolidation-related processes. Second, anodal tDCS may increase the excitatory-inhibitory (E-I) ratio (Barron et al., 2016) during training sessions, thereby promoting a more plastic state in the visual cortex (Bang et al., 2018; Shibata et al., 2017) and enhancing stimulus-specific learning gains in VPL following full practice. Future work is needed, integrating multimodal neuroimaging (e.g., fMRI-MRS fusion) to directly investigate functional reorganization and neurochemical plasticity in reactivation-versus repetition-based learning (Jia et al., 2023, 2024).

In sum, we propose reactivation-coupled brain stimulation as a combined intervention protocol for enhanced learning generalization at short training duration (i.e., reducing trial numbers by 80%), while maintaining learning gains. As memory reactivation mechanisms drive brain plasticity across multiple domains – including visual, motor, and mathematical learning – our reactivation-coupled anodal tDCS protocol may offer a translatable solution for clinical rehabilitation, enabling more efficient training with better generalization extending beyond the specific training conditions.

## Materials and Methods

### Participants

Night-six participants took part in the reactivation-based VPL experiment. Half completed an orientation detection task (main experiment; 21.71 ± 3.25 years old, 27 females) and half a motion detection task (control experiment; 21.38 ± 2.14 years old, 28 females). Within each task, participants were randomly assigned to the Reactivation/Anodal or the Reactivation/Sham group (N = 24 for each group). Based on a previous perceptual learning study using similar stimulation method (He et al., 2022), we conducted a prior independent t-test using the reported effect size (Cohen’s *d* = 0.9) in G^*^Power (Version 3.1) (Faul et al., 2007). This analysis indicated that 24 participants per group would provide power greater than 85% to detect the tDCS effect. Note that, this sample size was also comparable to prior tDCS studies on perceptual learning (Frangou et al., 2018; He et al., 2022; Jia et al., 2022). In addition, we recruited another forty-eight participants (22.00 ± 2.30 years old, 26 females) in the repetition-based VPL experiment, and randomly assigned them to the Full-Practice/Anodal or the Full-Practice/Sham group (N = 24 for each group). All participants were naïve to the purpose of the study, had normal or corrected-to-normal vision, and reported being right-handed. Written consent was obtained from all participants. The procedures used in this study were approved by the Ethics Committee at Department of Psychology and Behavioral Sciences, Zhejiang University (protocol number: 2022-061).

### Stimuli and apparatus

Gabor patches (Gaussian windowed sinusoidal gratings) were presented in the lower-right visual field at an eccentricity of 6.5° against a uniform gray background (∼35 cd/m ^2^). The Gabor stimuli had a diameter of 5°, random phase and spatial frequency of 1 cycle/degree. The Gaussian envelope had standard deviation of 2.5°. Noise patterns from sinusoidal luminance distributions were generated and superimposed on the Gabor patches at a specific signal-to-noise (S/N) ratio. For instance, a 20% S/N ratio indicates that the noise pattern replaced 80% of the pixels of the Gabor patch.

Random dot kinematograms (RDKs) were presented in an annular aperture located in the right visual field at 8°eccentricity. Each display contained 400 dots (0.1° diameter) moving at a speed of 10°/s. A specific proportion of dots moved coherently in one direction, while the rest moved randomly. When a dot moved out of the aperture, it was wrapped around to reappear from the opposite side along its motion direction.

The stimuli were generated using Psychtoolbox 3.0 (Brainard, 1997; Pelli, 1997) implemented in MATLAB (The MATHWORKS Inc., Natick, MA, USA). Stimuli were presented on a Dell Cathode-Ray Tube (CRT) monitor with the size of 40 × 30 cm ^2^, resolution of 1024 × 768 and a refresh rate of 60 Hz. Gamma correction was applied to the monitor. A chin-rest was used to stabilize participants’ head position and maintain the viewing distance at 72 cm.

### Experimental design and statistical analysis

Participants trained with the reactivation-based protocol completed five sessions in the following order: a pretest, an encoding session, two reactivation sessions, and a posttest. For participants trained with repetition-based protocol, the encoding and reactivation sessions were replaced with three standard full-practice training sessions (Figure 1A). All participants performed two-interval forced-choice orientation detection tasks throughout these sessions.

#### Orientation detection task

As shown in Figure 1B, each trial began with a central fixation cross (400 ms), followed by two sequential stimulus displays (50 ms each) separated by a 300 ms blank interval. One display contained a Gabor patch with specific S/N ratio, while the other contained pure noise (0% S/N ratio), with presentation order randomized across trials. Participants indicated which interval contained the Gabor patch via a keyboard press.

Participant’s performance in the task was measured using a 2-down 1-up staircase with 10 reversals converging at 70.7% performance. In each staircase run, the S/N started with 15% and adaptively changed with a step size of 0.05 log units. Each staircase run consisted of around 40 trials (1 – 2 mins). We calculated the thresholds as the geometric mean of the last six reversals. The reference orientation was set at 55° for the trained stimulus and 125° for the untrained stimuli, with these assignments counterbalanced across participants.

#### Behavioral test session

To stabilize fixation and familiarize participants with the task before tests, they first completed a 30-trial practice run (20% S/N ratio, above threshold). During practice run, auditory feedback was provided for incorrect responses. In both pretest and posttest, they completed four staircase runs of the orientation detection task (two runs per condition in random order). Detection thresholds were calculated by averaging the thresholds from the two runs per condition. No feedback was provided during tests.

#### Training session: Reactivation or Full-Practice

All participants were trained on an orientation detection task with fixed orientation and location throughout training sessions. Auditory feedback was provided for incorrect trials. The Full-Practice group (i.e., repetition-based VPL) completed three training sessions (16 staircase runs per session), while the Reactivation group performed 16 staircases runs on the encoding session, followed by two reactivation sessions, each consisting of three staircase runs. This design followed the protocol of a prior study (Bang et al., 2018), while also matched the duration of the online stimulation protocol (see *tDCS* section for details).

#### Behavioral data analysis

For each group and each orientation, we calculated a normalized learning gain index (NGI = [(Pre-test threshold – Post-test threshold) / ((Pre-test threshold + Post-test threshold) / 2)] × 100 %). Paired t-tests on NGI were used to compare performance between trained and untrained orientations within participants. To examine differences across multiple groups, we applied either independent t-tests or mixed ANOVAs on NGI. To quantify the amount of transfer effects, we calculated the NGI difference between the trained and untrained orientation (NGI difference = NGI_trained_ – NGI_untrained_). Lower NGI difference reflects more transfer, while higher NGI difference reflects greater specificity. A two-way independent-measures ANOVA (training protocol × stimulation condition) was applied on the NGI difference. To evaluate the strength of evidence for the lack of significant effects, we conducted parallel Bayesian analyses (Wagenmakers, 2007) using standard priors as implemented in JASP Version 0.17.1.0 (JASP Team, 2023).

#### Control experiment: motion detection task

To examine the robustness of the generalizable learning effect induced by reactivation-coupled brain stimulation, we replicated the reactivation-based experiment with motion stimuli (RDKs). The behavioral task procedure was similar to those used in the orientation detection task, with either anodal or sham stimulation. On each trial, two sequential displays were presented: one contained a signal RDKs with a given motion coherence, and the other was a noise field with 0% coherence (Figure 2A). The reference direction was set at 60°for the trained stimulus and 300°for the untrained stimuli, with these assignments counterbalanced across participants. Within each staircase run, the initial motion coherence was set to 15% and was adjusted adaptively using a step size of 0.05 log units.

#### *Transcranial direct current stimulation* (tDCS)

tDCS was delivered using a battery-driven, constant current stimulator with a pair of rubber electrodes in a 5 × 7 cm ^2^ saline-soaked synthetic sponges. In the main experiments of orientation detection task, the anode electrode was placed over the visual cortex (O1, 10-20 system) with conductive cream, while in the control experiment of motion detection task, the anode electrode was placed approximately 3 cm above the mastoid–inion line and 5 cm left of the midline in the sagittal plane (left V5, Battaglini et al., 2017). The cathode electrodes was positioned at the vertex (Cz) across experiments. Stimulation parameters followed safety guidelines. For the anodal tDCS condition, a direct current with an intensity of 1.5 mA was applied for 20 minutes, with a 30 s fade in/out periods to minimize cutaneous sensations. We used online stimulation protocol (i.e., stimulation during training). In particular, the current flow was initiated 10 minutes before task onset (rest period) and 10 minutes during the task. For the sham condition, participants received a 30 s fade-in phase followed by a 30 s fade-out at the beginning and end of the stimulation run, with no active stimulation in between. This sham protocol has been reported to effectively keep participants blinded to the stimulation conditions (Gandiga et al., 2006).

## Acknowledgments

This work was supported by grants to: K.J. from the National Natural Science Foundation of China (32571225, 32300855) and the Non-profit Central Research Institute Fund of Chinese Academy of Medical Sciences (2023-PT310-01). M.G. from National Natural Science Foundation of China (32371087), Fundamental Research Funds for the Central University (226-2024-00118), National Science and Technology Innovation 2030 — Major Project 2021ZD0200409. M.W. from National Natural Science Foundation of China (52407261), the “Pioneer” and “Leading Goose” R&D Program of Zhejiang (2025C01137). Z.K. from the Wellcome Trust (205067/Z/16/Z, 221633/Z/20/Z). For the purpose of open access, the authors have applied for a CC BY public copyright license to any Author Accepted Manuscript version arising from this submission.

## Author contributions

Y.X., M.G., and K.J. conceived the project and designed the experiments. Y.X., Y.G., B.W., and K.J. performed the experiments. Y.X., M.W., S.Z, and K.J. developed the tDCS protocols. Y.X., M.G., and K.J. developed the analysis pipeline and analyzed the data. Y.X., K.Z., M.G., and K.J. wrote the manuscript.

